# Lead (Pb) exposure results in cell type specific changes in the mouse retina and optic nerve

**DOI:** 10.1101/2025.08.16.670567

**Authors:** Labony Khandokar, Luke L. Liu, Wei Zheng, Patrick C. Kerstein

## Abstract

Chronic exposure to lead (Pb) is known to cause deficits in neuronal function across the nervous system, including the visual nervous system. Visual deficits have been observed in both humans and rodent models following Pb exposure. However, how Pb exposure causes visual deficits is poorly understood. In this study, we evaluated the effects of Pb toxicity on the retina and optic nerve of the mouse visual nervous system. We used C57BL/6 adult mice of both sexes and divided them into one of three different exposure groups. Adult mice received daily oral gavage of 108mg/kg Na-acetate (control), 54mg/kg Pb-acetate (low dose), or 108mg/kg Pb-acetate (high dose) for 4 weeks. At the end of Pb exposure, whole blood, retina, and optic nerve samples were collected for Pb quantification by atomic absorption spectroscopy and tissue immunohistochemical analyses. Cell type specific markers were used to quantify changes in cell density of retinal ganglion cells (RGCs), oligodendrocytes (OLs), oligodendrocyte precursor cells (OPCs), and myelin structure. Following Pb exposure, we observed a small, but significant reduction in the cell density of RGCs in the retina. However, we found no significant changes in branch thickness or coverage of retinal vasculature following Pb exposure. In the optic nerve after Pb exposure, we found a significant reduction in the cell density of OLs and OPCs. Finally, using immunolabeling for Caspr and Nav1.6, we observed significant structural changes in nodes of Ranvier, suggesting a disruption in myelin structure. Our findings suggested that Pb toxicity may impair survival and maturation process of oligodendrocytes, changes in myelin structures, and potential demyelination of the optic nerve. These results provide the foundation for future investigations into the molecular mechanisms of Pb-dependent changes in myelination and visual nervous system function.

## 1. Introduction

Environmental and occupational exposures to toxic substances cause functional deficits in the visual nervous system that result in impaired vision and blindness. Losses in vision can arise from specific exposures to chemicals and heavy metals (Fox, 2015; Saeed and Eldweik, 2025; Tshala-Katumbay et al., 2015). Exposure to the heavy metal lead (Pb), commonly found in drinking water, household products, and industrial byproducts, has been documented to cause structural and functional changes in several areas of the visual nervous system, including the retina, optic nerve, and visual cortex (Ebrahimi et al., 2024; Rosen et al., 2017). Despite the known effects of Pb on brain function controlling cognition and behaviors, the direct effects of Pb on the vision are poorly defined. Several studies have demonstrated elevation of Pb in the blood will led to Pb accumulation in the eye (Shen et al., 2016). Our earlier human study reveals that Pb accumulates in the retina at concentrations that are 166 and 739 times higher than those in the aqueous and vitreous humors, respectively (Eichenbaum and Zheng, 2000). Pb toxicity arising from its accumulation in the retina causes several clinical manifestations, include amblyopia, optic neuritis, optic atrophy, scotomas, and overall reduced visual function (Aghdam et al., 2019; Baj et al., 2022). Based on animal studies, Pb exposure may cause these symptoms through structural changes in the retina due to retinal neuron apoptosis, mitochondrial dysfunction, and changes in synaptic signaling (Gudadhe et al., 2024; Han et al., 2021; Jomova et al., 2025). In addition, studies in both human and animals during Pb exposure, have reported changes in physiological light responses by electroretinogram measurements from the retina and visual evoked potentials measured from the visual cortex (Sobieniecki et al., 2015).

Many visual disorders arise from the neurodegeneration of retinal photoreceptors or retinal ganglion cells (Telias et al., 2020; La Morgia et al., 2017). However, most studies in retinal toxicology, have focused their analyses on the dysfunction and cell death of photoreceptors. For example, in adult mice, Pb exposure induces apoptosis of photoreceptors through changes in mitochondria structure and function via the Bcl-X signaling pathway (Perkins et al., 2012). In addition, Pb exposure during retinal development also reduces the total number of photoreceptors; the mechanism of photoreceptor loss may be through reduced neurogenesis and differentiation rather than cell death (Fox et al., 2008). However, while most studies have focused on the effects on the retina through photoreceptors, the impact of Pb exposure on retinal ganglion cells and the optic nerve remains elusive.

Retinal ganglion cells (RGCs) play a vital role in transmitting information from the eye to the brain through the optic nerve. Despite the critical role RGCs plays in visual function, they are often susceptible to neurodegeneration and cell death in glaucoma, diabetic retinopathy, and several other optic neuropathies (Wang et al., 2025; Wong and Benowitz, 2022). These optic neuropathies have several etiologies but may arise from exposure to heavy metals, such as Pb (Baj et al., 2022; Saeed and Eldweik, 2025). For example, in humans, elevation of bone Pb levels correlates with the incidence of glaucoma (Wang et al. 2018). Structural changes have also been observed in the retina as the nerve fiber layer (containing RGC axons) is thinner in humans with higher blood Pb levels and therefore suggest a loss of RGCs or their axons (Aschner et al., 2024; Ekinci et al., 2014; Ebrahimi et al., 2024). Furthermore, childhood Pb exposure corresponds to changes in visual function, such as visually evoked potentials that test the connectivity between the eye and the brain (Altmann et al., 1998; Sobieniecki et al., 2015). Despite the association between Pb levels and optic nerve disorders associated with RGCs, the role of RGC death or degeneration has not been studied in animal and cell culture models. Therefore, the lack of these studies highlights the need for more animal models studying Pb exposure on RGC function and survival. These studies are necessary for our understanding of the underlying mechanism of Pb exposure on visual function.

When investigating optic neuropathies, it is important to address the other cell types that populate the optic nerve, such as oligodendrocytes, the myelinating cells of the central nervous system. Vision loss is a common symptom of demyelinating disorders, such as multiple sclerosis or neuromyelitis optica, highlighting the importance for myelination in normal vision (Jasse et al., 2013). Several studies have linked Pb exposure with reduced myelination levels in the central nervous system. For example, in human studies, higher blood Pb levels are correlated with reduced white matter tracts in the brain suggesting Pb may disrupt function through demyelination (Brubaker et al. 2009; Reuben et al. 2020). Similarly, following Pb exposure, both *in vitro* and *in vivo* studies in rodents have demonstrated a loss of myelin and oligodendrocytes in the brain (Deng et al. 2001; Liu et al. 2024). Within the visual nervous system, Pb-induced vision loss may involve a myelin-dependent process. Following Pb exposure, the loss of myelination in the optic nerve decreases nerve conduction velocity and connectivity between the eye and brain (Ethier et al. 2012). Similar functional deficits have been observed when oligodendrocytes were selectively ablated from the optic nerve (Balraj et al. 2023). While these findings highlight the role of myelin and oligodendrocytes in Pb-induced neurotoxicity, it is unknown whether similar mechanisms occur in the optic nerve. Further research is needed to understand the cellular basis of visual deficits observed following Pb exposure.

In this study, we aimed at exploration of Pb toxicity on retinal ganglion cells and oligodendrocytes in the retina and optic nerve following sub-chronic exposure to Pb in adult mice. After four weeks of Pb exposure, we found a small but significant decrease in the total number of retinal ganglion cells. In the optic nerve, we observed a decrease in both oligodendrocyte precursor cells and oligodendrocytes. Furthermore, we observed structural changes in the organization of nodes of Ranvier suggesting a reduction in the myelination of RGC axons. Finally, our data suggest Pb exposure induces cell death of retinal ganglion cells through the demyelination and loss of oligodendrocytes in the optic nerve.

## 2. Materials and methods

### 2.1 Animals and Lead (Pb) Exposure

For all experiments in this study, we used eight-week-old adult C57BL/6 mice (Jackson Laboratory). The mice were randomized into three groups, housed within a temperature-controlled environment, and adhered to a 12-hour light/dark cycle. Each group consisted of three males and three females. Exposures were administered once daily via oral gavage 5 days/week for 4 weeks. The control group received an oral gavage of 100 μL of 66 mM sodium acetate (108 mg/kg), while the two experimental groups were exposed to “low” 33mM (54 mg/kg) or “high” 66 mM (108 mg/kg) Pb acetate (Figure 1A). All procedures related to the care and handling of the animals were approved by the Purdue University Institutional Animal and Use Committee and followed the National Institute of Health’s *Guide for the Care and Use of Laboratory Animals*. All animal experiments complied with the ARRIVE (Animal Research: Reporting of In Vivo Experiments) guidelines.

**Figure 1.**
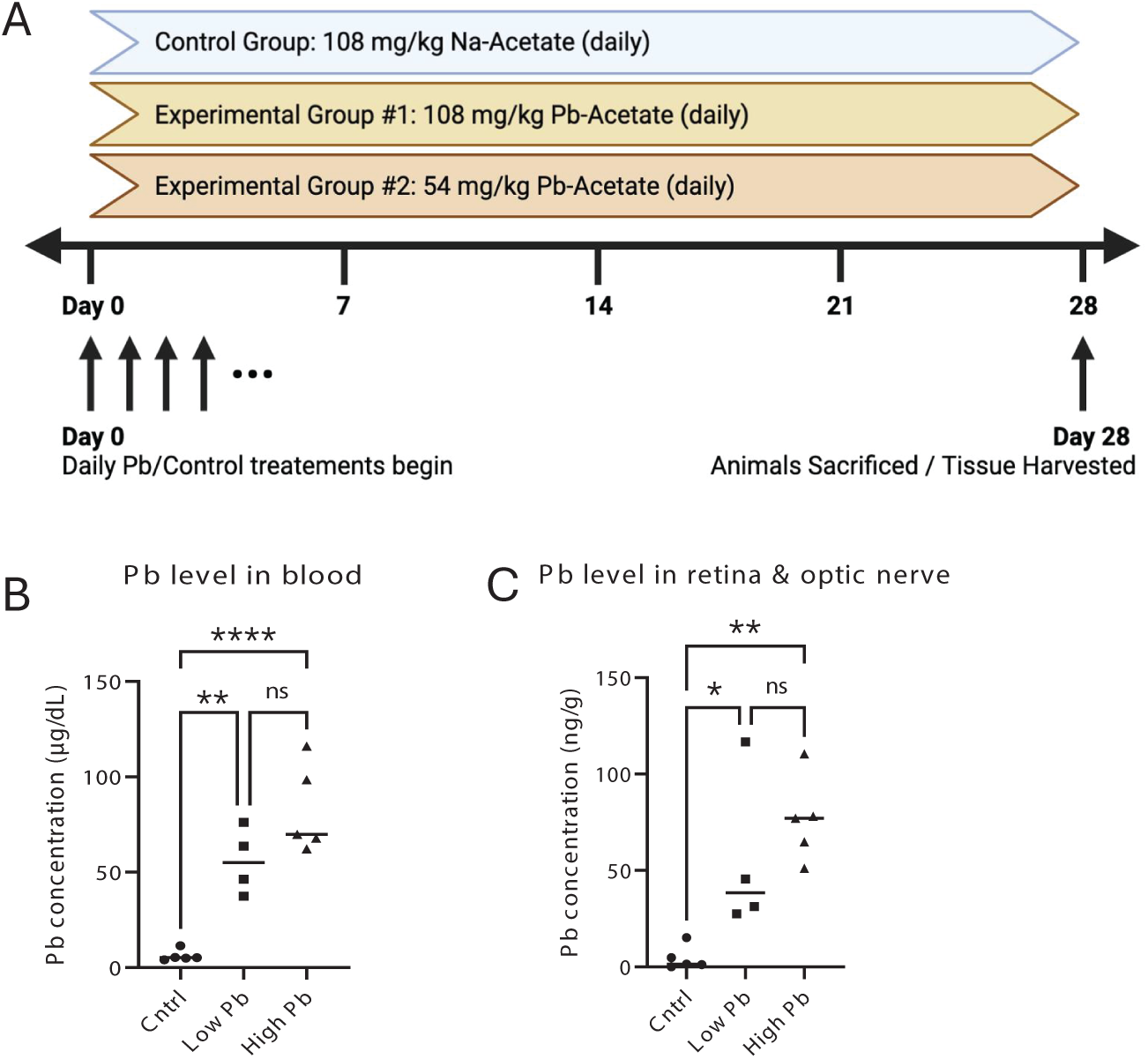
Chronic Pb exposure elevates Pb concentration in the whole blood, retina, and optic nerve of adult mice. (**A**) A schematic depicts the daily oral gavages mice received over four weeks. Mice were separated into 3 exposure groups that received either 108 mg/kg sodium acetate (control), 54 mg/kg Pb acetate (Low Pb), or 108 mg/kg Pb acetate (High Pb). (**B-C**) Quantification of Pb levels by atomic absorption spectroscopy in (**B**) whole blood or (**C**) retina and optic nerve samples from each exposure group. Data presented as mean ± SEM. n= 4-6 mice per exposure group. ns = not significant; *p < 0.05; **p < 0.01; and ****p < 0.0001 using a One-way ANOVA with a Tukey’s multiple comparison post-hoc test.

### 2.2 Tissue Collection and Preparation

Within 24hrs of the final exposure, mice were euthanized using carbon dioxide followed by cervical dislocation. The eyes were removed and placed in 4% paraformaldehyde (PFA) for 30 mins at room temperature. Next, the eyes were washed twice in 1X PBS for 30 mins. After fixation, retinas and optic nerves were dissected free from the eye for cryosections or flat-mounts. For cryosections, optic nerves were cryoprotected in 15% sucrose solution for 2 hrs at 4°C, frozen in cryo-molds containing optimal cutting temperature (O.C.T.) media and stored at -80°C. Retina and optic nerves were sectioned on a cryostat at 20µm and collected on a glass microscope slide (Fisher Superfrost Plus). For retinal flat mounts, after fixation, retinas were removed from the eye. Finally, retinas were flattened by making 3-4 equally spaced incisions around the edges of the retina in preparation for immunohistochemistry and mounting on glass slides for fluorescence microscopy.

### 2.3 Blood and Tissue Pb Quantification

Within 24 hrs of the last dose, mice were sacrificed, and blood samples were collected from the inferior vena cava into 100 µL of 0.5M EDTA. Tissue samples (retina and optic nerves) were collected as described above. Blood and tissue samples were measured for volume and weight, respectively. Both blood and tissue samples were digested at 120 °C for 16 hours in a closed Teflon container with 65% metal-free HNO_3_. Pb levels were analyzed by graphite furnace atomic absorption spectrometry (GFAAS) using a Perkin Elmer 4100 ZL spectrometer (Perkin Elmer, Warsaw, Poland). All diluted samples had Pb concentration within the linear range (0 to 20 μg/L) of this AAS method. Pb standard curves generated using Agilent’s Pb standard solution were plotted and monitored to consistently achieving an R2 value exceeding 99.5%, as described previously (Liu et al. 2024).

### 2.4 Immunohistochemistry

For immunolabeling, slide-mounted optic nerve sections were washed with 1X PBS for 10 mins and then blocked with normal donkey serum (NDS) with 0.2% Triton X-100 for 30 mins. Retinal sections were then incubated with primary antibodies in NDS blocking buffer overnight at 4°C. The primary antibodies and their dilutions used in this study are listed in Table 1. Next, sections were washed 3 times in NDS blocking buffer for 10 mins and incubated in donkey secondary antibodies in blocking buffer for 2 hours at room temperature. Retinal sections were washed three times in PBS for 10 mins with either Hoechst or DAPI (1:5000) stain included in the first wash step. The tissue was mounted in Fluoromount-G (Southern Biotech) and cover-slipped for imaging.

**Table 1:**
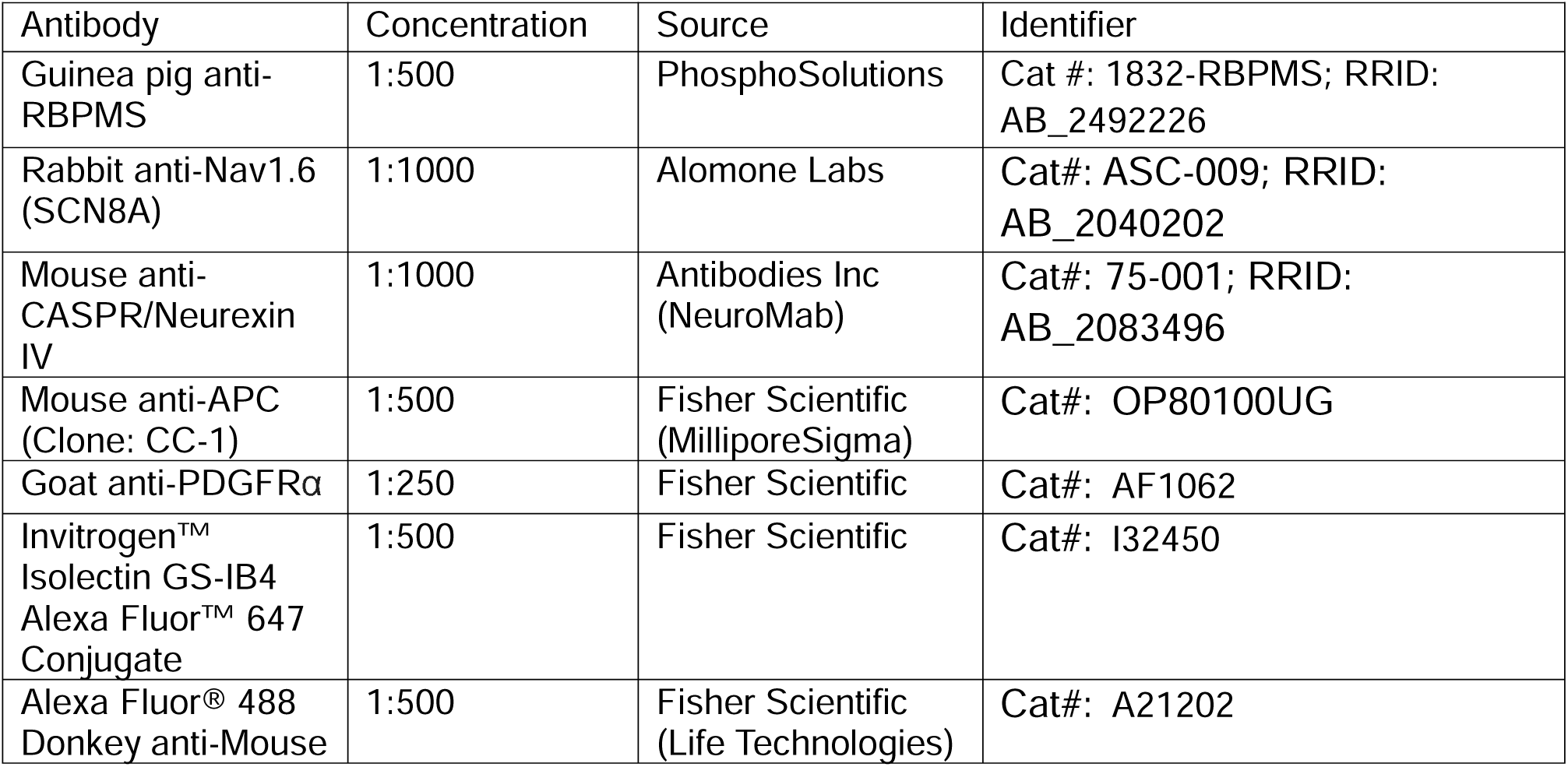
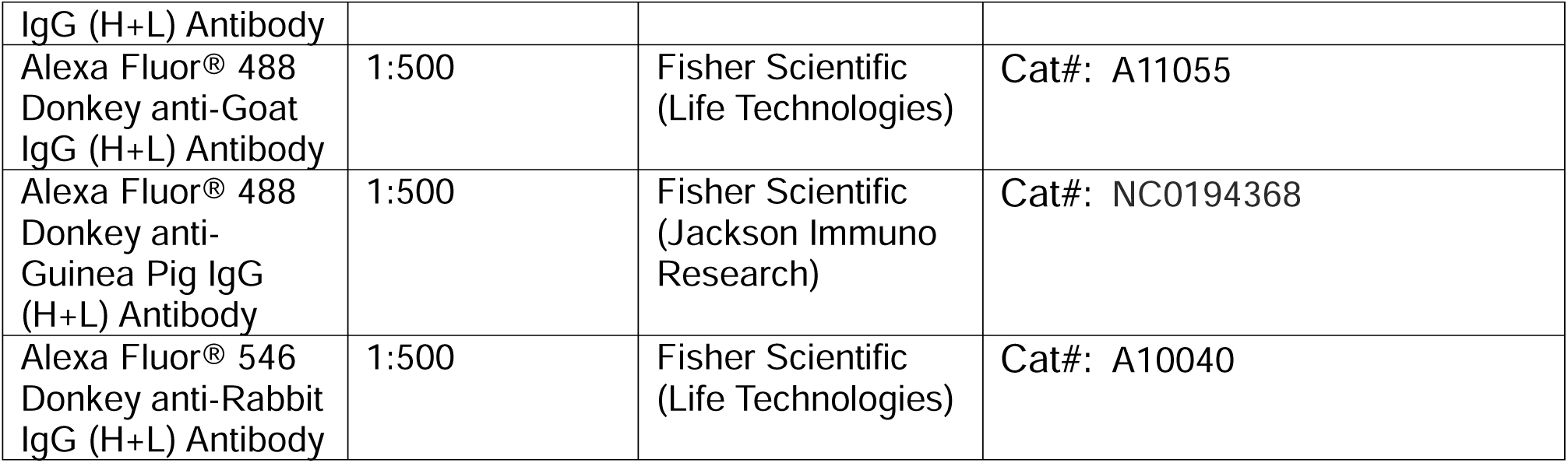

For flat mounts, retinas were washed 3 times and blocked in 2% NDS and 0.2% Triton X-100 for 30 mins at room temperature. Retinas were incubated with primary antibodies in blocking solution for 3-7 days at 4°C with gentle agitation. After incubation with primary antibody, the retinas were washed 3 times with blocking solution for 30 mins. Next, retinas were incubated with secondary antibody in blocking solution overnight at 4°C. For retinal vasculature labeling, retinal flat-mounts were incubated with 10 µg/mL Isolectin-B4 conjugated to Alexa Fluor 647 (Invitrogen) overnight at 4°C. Finally, retinas were then washed three times in PBS for 30 mins with either Hoechst or DAPI (1:5000) stain included in the first wash step. Tissue was mounted on a slide with Fluoromount-G and cover-slipped for imaging.

### 2.5 Fluorescence image acquisition

All retinal and optic nerves were imaged on a Zeiss LSM 900 laser-scanning confocal microscope using Zeiss Zen software. For optic nerve sections, immunolabeled cells were capture using a 20X objective. Node of Ranvier structures were captured using a 63X Oil objective. For retinal flat mounts, a 20X objective was used for cell density measurements and whole retinas were captured by stitching together images captured by a 10X objective. For all immunolabeling experiments, 6 different images were acquired per animal for off-line analysis.

### 2.6 Fluorescence image analysis

All images were analyzed off-line using the Fiji software (Schindelin et al., 2012). Cell density measurements in retinas and optic nerves for RGCs (RBPMS), oligodendrocytes (CC1), and oligodendrocyte precursor cells (PDGFRα) were quantified from a 250×250µm area. For RGC density, 4 measurements were made across the retina from regions at least 500µm from the optic nerve head and the retina edge. For retinal vasculature branch measurements, primary and secondary branch measurements were made from retinal flat mounts. Primary branches were defined as branches originating from the optic nerve head and secondary branches were defined as branches originating from primary branches. For each retina, at least 6 measurements were made from the central regions of the retina. In the optic nerve, nodes of Ranvier were defined as two in-line CASPR-labeled puncta separated by a single Nav1.6-labeled puncta. The paranode region was measured as the width across two CASPR-labeled puncta. The node region was measured as the width between the two CASPR-labeled puncta. The Nav1.6 cluster region was measured as the diameter the Nav1.6 puncta. For node of Ranvier analysis, at least 15 measurements were made from 6 images from each optic nerve.

### 2.7 Statistical analysis

In these analyses, each experiment included measurements from a minimum of 4 retinas from at least 4 different animals. For all datasets, the measurements and variance were reported as mean ±SEM. Each dataset was first tested for a normality with a D’Agostino & Pearson test. For analysis between multiple groups (≥3 groups), we used both One-way and Two-way analysis of variance (ANOVA) with a Tukey’s multiple comparisons test (parametric) or a Kruskal-Wallis with a Dunn’s multiple comparison test (nonparametric). All statistical tests were performed using Prism 10 software (Graphpad Software, Inc.).

## 3. Results

### 3.1 Lead (Pb) exposure elevates Pb levels in the whole blood, retina, and optic nerve

To assess the effects of Pb exposure on the retina and optic nerve we treated mice with a sub-chronic Pb exposure. Adult mice received daily oral gavage of Na-Acetate (108 mg/kg) as control, a “low” dose of Pb-Acetate (54 mg/kg), or a “high” dose of Pb-Acetate (108 mg/kg) for 4 weeks (Figure 1A). This sub-chronic Pb exposure significantly increased blood lead levels (BLLs) in both treatment groups as compared to the control group (p < 0.0001; One-Way ANOVA with Tukey’s post hoc test). After 4 weeks of treatment, the mean BLLs in the control group were 6.16 μg/dL whereas mean BLLs in the low Pb group and high Pb group were 55.90 μg/dL and 82.84 μg/dL, respectively (Figure 1B).

The Pb exposure also led to a significant accumulation of the this metal in the retina and optic nerve in treated groups (p < 0.01; One-Way ANOVA with a Tukey’s post hoc test). The mean Pb concentrations in the retina and optic nerve of the low and high Pb-exposed animals were 55.3 ng/g and 76.3 ng/g, respectively, whereas the mean lead concentration of the control group was 4.5 ng/g (Figure 1C), about a 12-fold and 17-fold increase over controls, respectively.

### 3.2 Pb exposure decreases Retinal Ganglion Cell (RGC) density

A major characteristic of optic neuropathies is the reduction of cell density of RGCs due to cell death during the disease process (Carelli et al. 2017). Here, we aimed to determine if Pb exposure affects the cell density of RGCs in the retina. For this purpose, flat mount retinas were immunolabeled with RGC marker RBPMS to measure the density of RGCs following Pb exposure (Figure 2A-F). After 4 weeks of Pb exposure, we found a 15.5% reduction in the cell density of RBPMS+ cells in the retinas from the high Pb treated group (p < 0.01, One-way ANOVA with Tukey’s post hoc test) (Figure 2G). Similar to increase in Pb concentration in the retina (Figure 1B), we found a small, but significant reduction in RGC density in the low and high Pb treated groups compared to the control group (Fig. 2B, E, G). These data suggest that Pb exposure causes a loss of RGCs potentially through cell death over a 4-week period.

**Figure 2.**
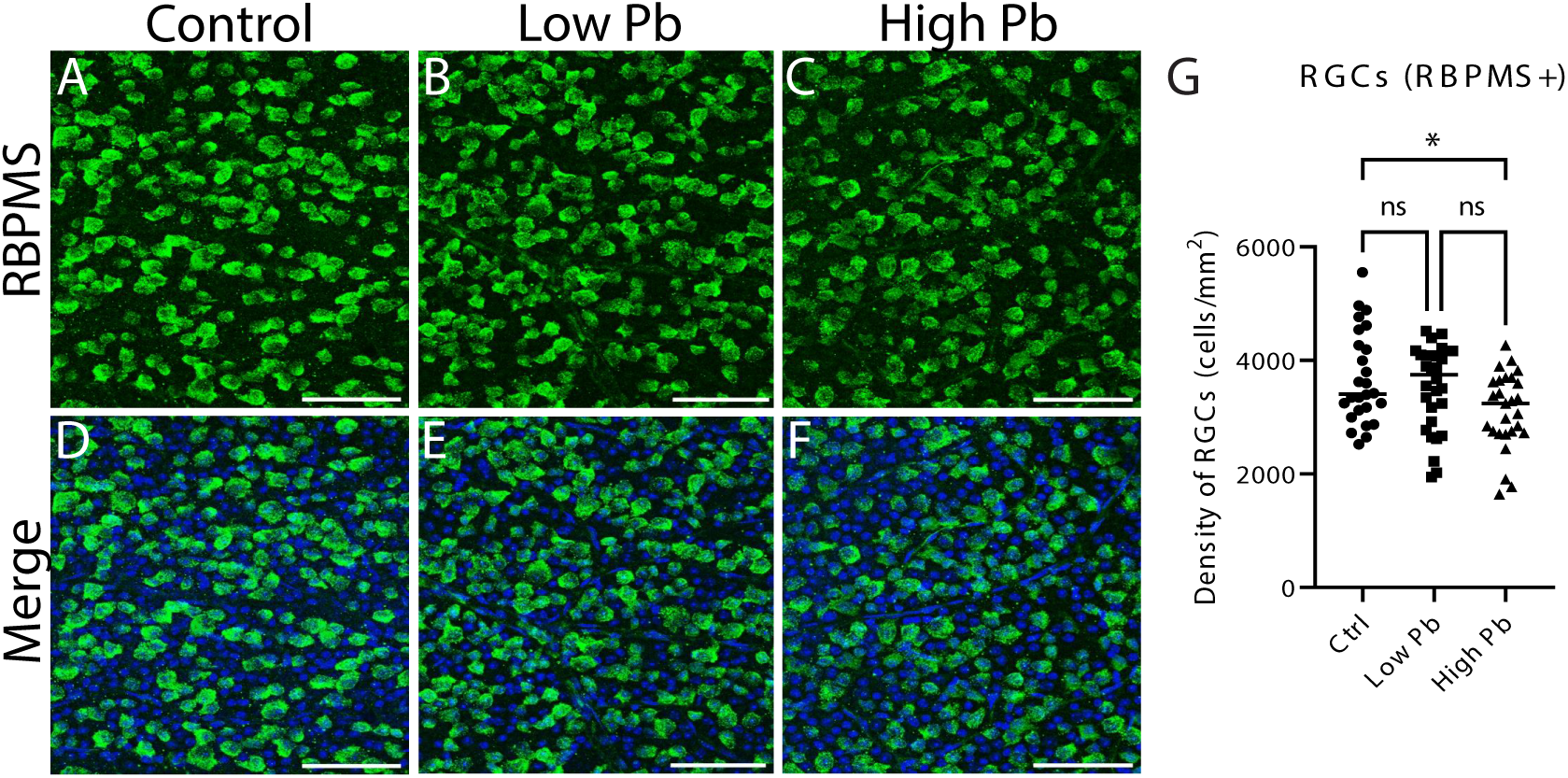
Retinal ganglion cells (RGCs) loss after Pb exposure. (**A-F**) Retinal flat mounts were immunolabeled with the RGC marker RBPMS (green) in (**A**) control, (**B**) low Pb, and (**C**) high Pb groups. (**D-F**) Images from A-C are merged with nuclear marker DAPI. (**G**) Quantification of RGC density from control, Low Pb, and High Pb treatment groups. Data presented are mean ± SEM. n=4 measurements per retina, 6 mice per exposure group. ns, not significant; *p<0.05; **p<0.01; One-way ANOVA with a Tukey’s multiple comparison post-hoc test. Scale bar, 80 µm in (**A-F**).

### 3.3 Pb exposure does not alter the retinal vasculature structure

Retinal injury and disease can lead to changes in retinal vasculature integrity and structure. Furthermore, several optic neuropathies, such as diabetic neuropathy, exhibit blood hemorrhages and neovascularization (Eggers, 2023). In this study, we wanted to investigate whether Pb exposure induces changes in the overall retinal vasculature structure and organization. To label the retinal vasculature, we stained retinal flat mounts with Isolectin B4 (IB4). We observed no changes in the overall vasculature coverage in the retina with Pb exposure (Figure 3 A, D). In addition, the retinal vasculature density in both central and peripheral retina remained similar in all exposure groups (Figure 3B-C, E-F). One of the explicit clinical manifestations in acute Pb poisoning is brain swelling, accompanied with herniation, ventricular compression, and petechial hemorrhages (Pentschew 1965; Smith et al., 1960; Goldstein et al., 1974). Evidence also shows a distinct cerebral endothelial damage along with the blood-brain barrier leakage following chronic Pb exposure (Wang et al., 2008). To further evaluate retinal vasculature structure, we quantified both primary and secondary branch thickness. Following Pb exposure, we did not observe a significant difference of mean blood vessel thickness between control and Pb-exposed groups (Figure 3G-H). Unlike previous studies in the brain, retina vasculature structure remains relatively normal following Pb exposure.

**Figure 3.**
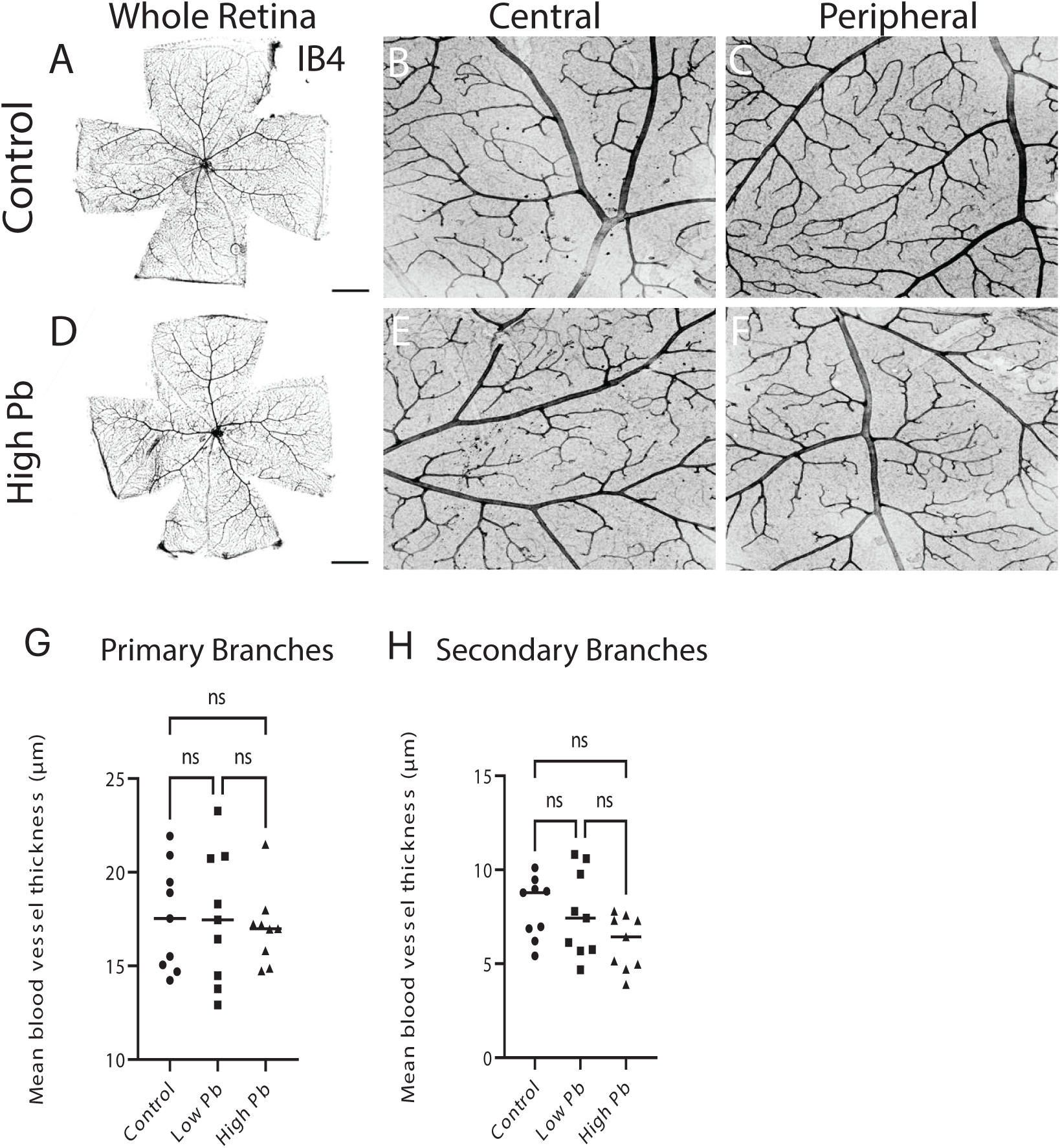
Retinal vasculature structure remains unchanged after Pb exposure. (**A-F**) (**A-F**) Retinal flat-mounts were labeled with blood vasculature marker Isolectin-B4 (IB4) in the (**A-C**) control and (**D-E**) high Pb exposure groups. (**B-F**) Magnified images were presented from (**B, E**) central and (**C, F**) peripheral region of retinal flat mounts from (**B-C**) control and (**E-F**) high Pb exposure groups. (**G-H**) Quantification of mean blood vessel thickness for (**G**) primary and (**H**) secondary branches. Data presented as mean ± SEM. n≥9 total measurements from at least 6 mice per exposure group. ns, not significant. Statistical analysis used a One-way ANOVA with Tukey’s multiple comparison post-hoc test. Scale bar, 400 μm in (**A, D**) and 50μm in (**B**).

### 3.4 Pb exposure decreases the cell density of oligodendrocyte precursor cells and oligodendrocytes

Several studies have suggested Pb exposure may also lead to a reduction in myelination along axon tracts of the central nervous system (Rai et al. 2013; Liu et al. 2024). Demyelination is often the result of loss of myelinating cells, i.e., oligodendrocytes (OLs) and oligodendrocyte precursor cells (OPCs). To evaluate the effect of Pb exposure on the myelinating cells of the optic nerve, we examined the cellular density of OPCs and OLs. We quantified the cell density by immunolabeling with each cell type with either the OPC marker PDGFRα or the OL marker CC1. We observed a significant lower cell density of OLs in both Pb-exposed groups (Figure 4B, D) compared to the controls (Figure 4A, C). The mean OL density was reduced by 28.6% and 40.2% compared to control mice in low and high Pb-exposed groups, respectively (p< 0.0001; One-Way ANOVA with a Tukey’s post hoc test) (Figure 4I). Similarly, the cell density of OPCs cells was significantly reduced in both Pb-exposed groups compared to the control group (Figure 4E-H). The mean cell density of OPCs was reduced by 32.4% AND 36.6% compared to control mice in the low and high Pb-exposed mice, respectively (p< 0.001; One-Way ANOVA with a Tukey’s post hoc test). These data show that Pb exposure reduces the cell density of OPCs and OLs in the optic nerve possibly leading to visual deficits through demyelination of the RGC axons.

**Figure 4.**
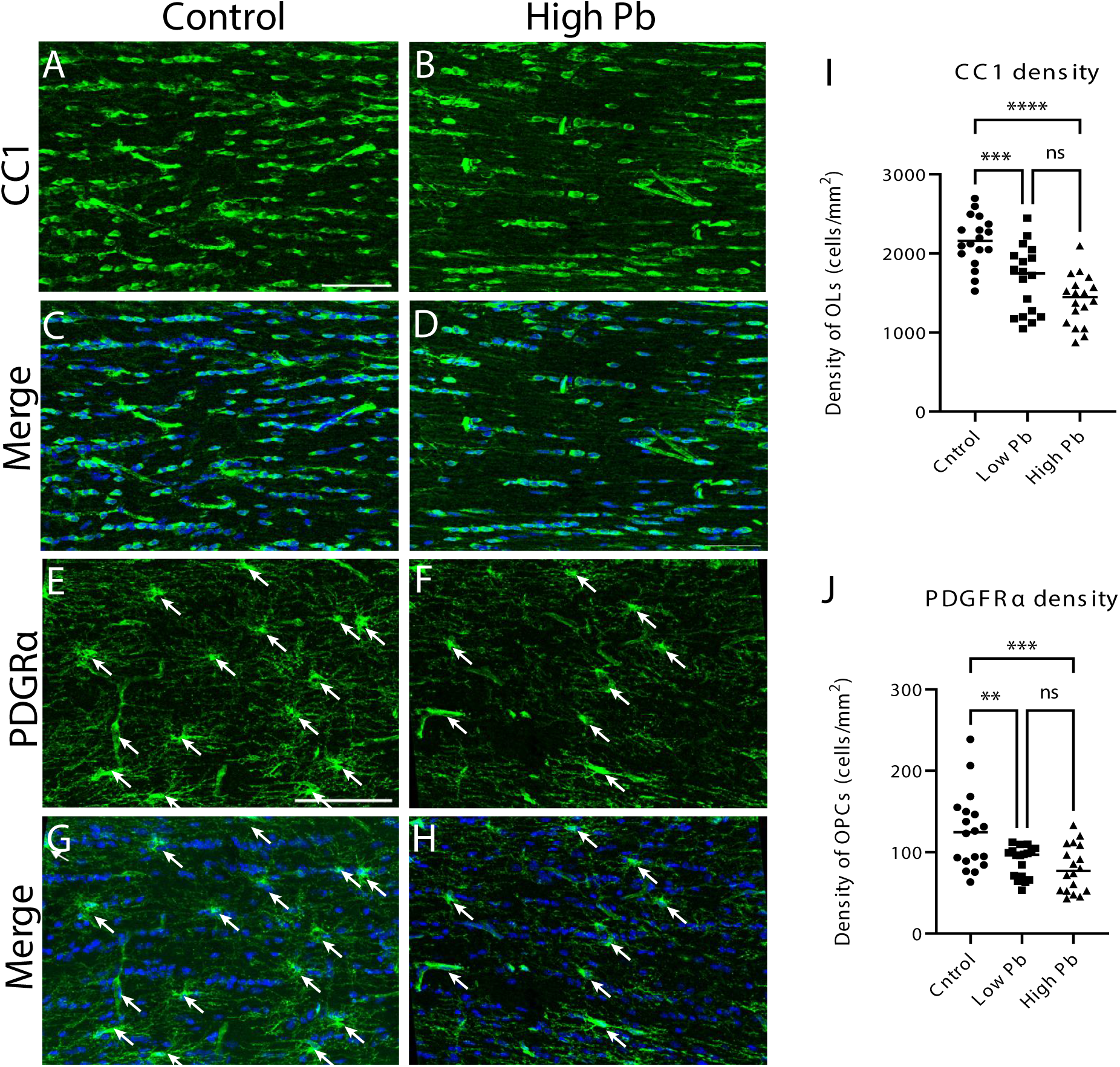
Pb exposure reduces the cell density of oligodendrocytes (OLs) and oligodendrocytes precursor cells (OPCs) in the optic nerve. (**A-D**) Optic nerve sections are immunolabeled with OL marker CC1 in (**A, C**) control and (**B, D**) high Pb exposed animals. (**C-D**) Images from A-B are merged with nuclei marker DAPI. (**I**) Quantification of OL (CC1) cell density across exposure groups. (**E-H**) Optic nerve sections are immunolabeled with oligodendrocyte precursor cell marker PDGFRIZl in (**E, G**) control and (**F, H**) high Pb exposed animals. (**G-H**) Images from A-B are merged with nuclei marker DAPI. (**E-H**) White arrows indicate cell bodies of PDGFRIZl-positive OPCs. (**J**) Quantification of OPC (PDGFRIZl) cell density across Pb exposure groups. n=6 measurements from 6 mice per exposure group. Data presented as mean ± SEM. ns, not significant, \*\**p* < 0.01, \*\*\**p* < 0.001 and \*\*\*\**p* < 0.0001 using a One-way ANOVA with Tukey’s multiple comparison post-hoc test. Scale bar, 80 μm in (**A, E**).

### 3.5 Altered node of Ranvier organization following Pb exposure

To further evaluate the effect of Pb exposure on myelination of the optic nerve, we used immunohistochemistry to label distinct structures of the myelin sheath and nodes of Ranvier. Immunomarkers, CASPR and Nav1.6, were used to label the paranode region and sodium channel clusters within the node region, respectively (Figure 5A-H). Changes in these structures has been associated with a functional vision loss in eye disorders (Wu et al. 2023). In optic nerve sections, based on CASPR-labeling, we did not observe any changes in the paranode length between the Pb-exposed and control mice (Fig. 5I). However, we observed an irregular distribution of Nav1.6 and an increase node length in the high Pb group compared to the control group (Figure 5J-K). The node length significantly increased by 58.6% and 67.7% compared to control mice in the low and high Pb-exposed mice, respectively (p< 0.0001; One-way ANOVA with a Tukey’s post-hoc test) (Fig 5J). Furthermore, following Pb exposure, the width of the Nav1.6 cluster in the node was significantly reduced by 29.3% in the high Pb-exposed mice compared to controls (p< 0.01; One-way ANOVA with a Tukey’s post hoc test) (Fig. 5K). Overall, these data confirm that Pb exposure alters node of Ranvier organization, suggesting the shorter myelin sheath length and reduced electrical signaling along the axons of the optic nerve.

**Figure 5.**
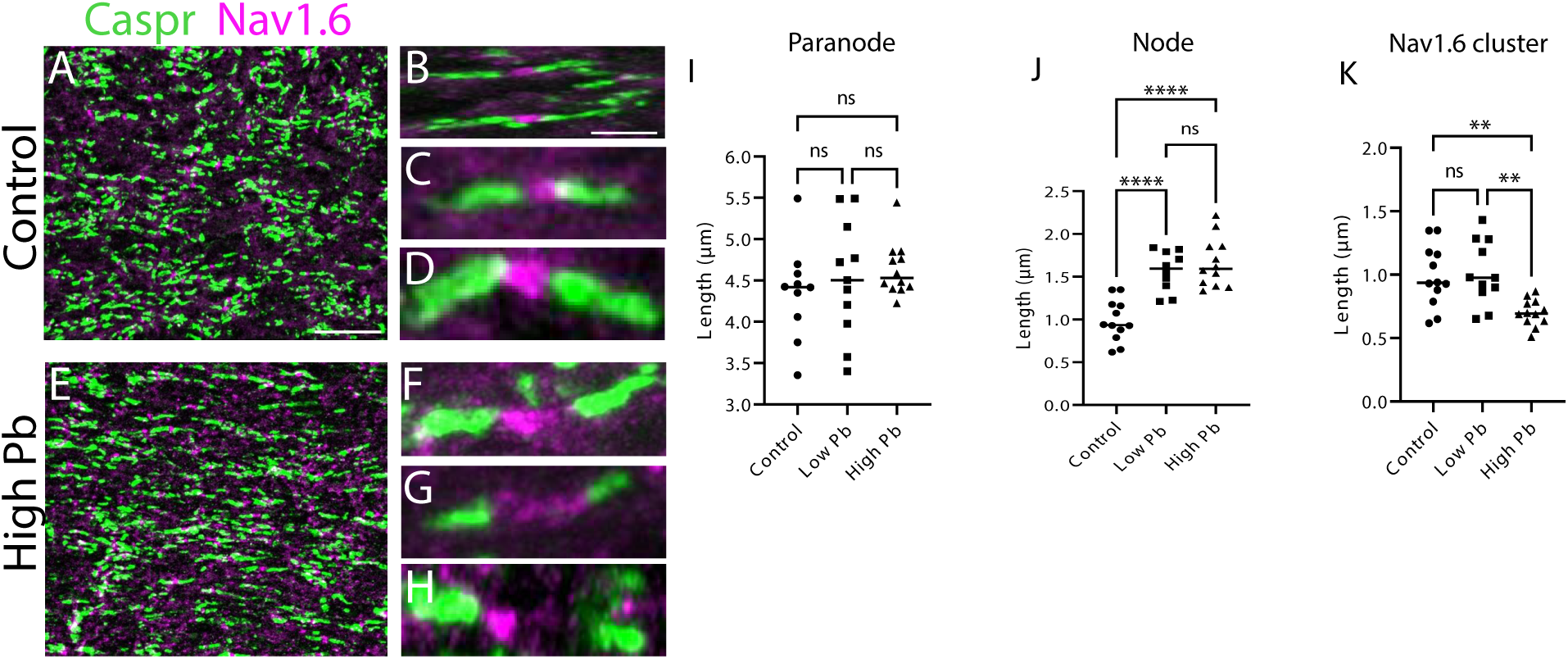
Pb exposure disrupts node of Ranvier structure. (**A-H**) Optic nerve cryosections were immunolabeled with paranode marker Caspr (green) and node marker Nav1.6 (magenta) antibodies in the (**A-D**) control and (**E-H**) high Pb exposure groups. (**B-D, F-H**) Magnified images showing examples of single nodes of Ranvier in the (**B-D**) control and (**F-H**) high Pb exposure groups. Quantification of (**I**) Paranode length, (**J**) node length, and (**K**) Nav1.6 cluster width across exposure groups. Data presented as mean ± SEM. n=15 measurements from at least 6 animals per exposure group. ns, not significant, \*\**p* < 0.01, \*\*\**p* < 0.001 and \*\*\*\**p* < 0.0001 using a One-way ANOVA with a Tukey’s multiple comparison post-hoc test. Scale bar: 80 μm in (**A**) and 10 μm in (**B**).

## 4. Discussion

In the central nervous system, Pb exposure causes demyelination and neuroinflammation in several distinct regions of the brain (Liu et al., 2024; Chibowska et al., 2016; Dehghanifiroozabadi et al., 2019). Visual impairments can also occur with Pb exposure suggesting Pb toxicity may have similar effects in the visual nervous system (Sobieniecki et al., 2015). In the current study, we evaluated the effect of sub-chronic Pb exposure on the retina and optic nerve of the mouse visual nervous system. We found that Pb accumulates in the retina and optic nerve of the eye (Figure 1). After 4 weeks Pb exposure, we observed a small, but significant, loss of retinal ganglion cells (RGCs) in the retina (Figure 2). Furthermore, in the optic nerve, Pb exposure caused both a significant loss of OLs and OPCs and altered myelin organization (Figure 4-5). Finally, while Pb exposure has previously been shown to cause brain vasculature thinning (Wang et al. 2008), in this study, we did not observe any structural changes in retinal vasculature (Figure 3). These data provide first-hand evidence that Pb exposure causes an optic neuropathy that may potentially lead to permanent RGC loss in the retina.

How does Pb toxicity affect the survival of RGCs, OLs, and OPCs? Several molecular and cellular mechanisms may explain Pb toxicity on the visual nervous system. Central to this question are RGCs that have both a high metabolic activity and a high energy demand, making them more vulnerable to oxidative stress, excitotoxicity, mitochondrial dysfunction, and inflammatory insults (HollaLJnder et al., 1995). RGC loss resulting from Pb toxicity may occur through one of a few mechanisms: 1) the loss of supporting and myelinating cells, 2) direct disruption of mitochondria in RGCs, and 3) the generation of reactive glial cells through neuroinflammation pathways.

First, Pb toxicity may affect RGC function and survival indirectly through the disruption of myelination along RGC axons in the optic nerve. In this study, we observed a loss of both OLs and OPCs following Pb exposure. This is further supported by our recent study (Liu et al., 2024) that demonstrated that chronic Pb exposure disrupts oligodendrogenesis in the brain’s regenerative niche—the subventricular zone. This disruption results in distinct demyelination in the corpus callosum detected by both immunohistochemistry and MRI assessment. Demyelination and the loss of myelinating cells has been shown to induce RGC cell death. For example, in animal models, selective ablation of OLs results in RGC dysfunction and eventual cell death, therefore suggesting that both myelination and OL function are essential to RGC survival (Balraj et al. 2023). The loss of OLs leading to RGC dysfunction has also been observed in human patients and animal models of disease. In the demyelinating disorder multiple sclerosis, the immune response triggers a chain of pathological pathways that cause injury of both OLs and OPCs leading to demyelination and eventual axonal damage and degeneration (Lassman and van Horssen 2015; Errea et al., 2015; Dutta et al., 2006; Dulamea, A. O. 2017). Similarly, in models of diabetic retinopathy, impairment of OPC and OL differentiation and survival reduces RGC axon number and diameter leading to visual impairments (Fernandez et al., 2012; Zhou and Chen, 2023; Wu et al., 2023). Previous studies have also shown that Pb toxicity effects the survival and function of OPCs and OLs in central nervous system myelination. For example, in cell culture, Pb toxicity inhibited the maturation and survival of OPCs (Deng et al. 2001; Ma et al. 2015). These studies, in agreement with the current study, suggests Pb toxicity disrupts the survival and potential maturation of OPCs and OLs leading to reduced myelination along axon tracts of the central nervous system. The loss of OPCs and OLs may lead to the subsequent degeneration and cell death of RGCs.

Second, both environmental and genetic disruption of mitochondria function can lead to the degeneration and loss of RGCs. In both human patients and animal models, this RGC dysfunction and death can arise from mitochondrial abnormalities associated with oxidative stress, reduced mitochondrial respiratory activity, and alterations in mitochondrial DNA (Abu Amero et al., 2006; Almasieh et al., 2012). In addition, alterations in proteins involved in mitochondrial function leads to hereditary optic neuropathies, such as Leber’s hereditary optic neuropathy and dominant optic atrophy, that are characterized by the degeneration of RGC axons (Baderna et al., 2020; Yang et al. 2024). Mitochondrial dysfunction has long been known to be associated with Pb toxicity; it may act as a direct mechanism of RGC dysfunction. Pb exposure has been shown to impair mitochondrial energy metabolism by reducing ATP production and oxygen consumption, along with decreasing the activities of essential enzymes (Han et al., 2021). Pb toxicity may target these mitochondrial processes to directly affect the survival of RGCs in optic neuropathies.

Third, the neuronal and OL loss during Pb toxicity may occur through the activation of other glial cell types, such as astrocytes and microglia. Briefly, astrocytes and microglia are essential glial cells in the central nervous system that play roles in maintaining homeostasis and regulating the immune response in various pathological conditions (Matejuk and Ransohoff, 2020). One potential mechanism of Pb toxicity in the optic nerve is through the activation of astrocytes and microglia. In recent studies, Pb toxicity induces a neuroinflammation response through the activation of both astrocytes and microglia (Gąssowska-Dobrowolska et al., 2023). Specifically, Pb toxicity has several effects on astrocytes, such as generation of reactive astrocytes, gene expression changes observed with injury or cell death, and responses to cytokines released by microglia (Kiray et al., 2016; Ponath et al., 2018). These cytokines have been previously demonstrated to cause neurodegeneration and cell death of OLs and OPCs (Allaman et al., 2011; Kiray et al., 2016; Ludwin, et al., 2016; Miljkovic et al., 2011; Ponath et al., 2018). In addition to astrocytes, Pb exposure induces microglial activation by causing the release of nitric oxide, glutamate, or the expression of pro-inflammatory cytokines, such as IL-1β, IL-6, and TNFα (Liu et al., 2012). Secretion of these factors reduce both OPC differentiation and OL maturation (Kalafatakis and Karagogeos, 2021; Li et al., 2008; Pang et al., 2010). Finally, microglia-astrocyte crosstalk can also lead to OL cell death. For example, activated microglia also facilitates the activation of astrocytes that inhibit OPC differentiation and induces OL cell death (Brambilla, 2019; Colombo and Farina, 2016; Kalafatakis and Karagogeos, 2021; Liddelow et al., 2017). Therefore, in future studies, we will need to explore the role of glial cells in the optic nerve, especially microglial and astrocytes in the neuroinflammation process induced by Pb exposure in relation to OL and OPC injury.

Finally, Pb appears to have a unique affinity to cerebral endothelial cells, as it accumulates in much greater concentrations in brain endothelial cells than other brain cell types (Struzynska et al., 1997; Toews et al., 1978). Surprisingly, in the present study we did not observe any significant structural damage in retinal blood vessels after Pb exposure. This could be due to the sensitivity of the retinal blood vessels that is different than the cerebral blood vessels. It could be also due to the species difference between rodents and humans.

In summary, adult Pb exposure reduces RGC density, impairs OL maturation or survival, alters myelin structure, and causes potential demyelination of the optic nerve. These data suggest that OPCs and OLs may be the cellular target of Pb toxicity in the visual nervous system. Mechanistically, the loss of OLs and demyelination in the optic nerve may result from Pb-induced oxidative stress, mitochondrial dysfunction, reactive gliosis of microglia and astrocytes; however, these mechanisms need to be further explored in future studies. This study provides the foundation for future investigations into the molecular mechanisms of Pb-dependent changes in myelination and visual nervous system function.

## Acknowledgements

The authors would like to thank the members of Kerstein and Zheng labs for their comments on this manuscript. The research in this manuscript was funded by the Showalter Research Trust (to P.C.K.), lab start-up funds from Purdue University (to P.C.K.), and an NIH/NIEHS R01 ES027078 grant (to W.Z.).

## Notes

### Competing Interest Statement

The authors have declared no competing interest.

